# Performance Evaluation and Selection of Improved *Tef* (*Eragrostis tef* L) varieties at Main Campus Site Hadiya Zone, Southern Ethiopia

**DOI:** 10.1101/2020.08.28.267583

**Authors:** Dargicho Dutamo, Ermias Assefa, Muluneh Menamo

**Affiliations:** College of Agricultural science, Wachemo University, Ethiopia. P.O.Box: 667; Ethiopian biotechnology institute, Addis Ababa, Ethiopia

**Keywords:** Adaptability, Grain, Yield, Tef, Varieties

## Abstract

Tef [Eragrostis tef (Zucc.) Trotter] is a cereal crop resilient to adverse climatic and soil conditions, and possessing desirable storage properties. It is, a tetraploid with 40 chromosomes (2n = 4x = 40), belongs to the family Poaceae and, together with finger millet (Eleusine coracana Gareth.), to the subfamily Chloridoideae. It was originated and domesticated in Ethiopia. The experiment was conducted to identify, select and recommend adaptable, high yielding, Insect pest and disease resistant twelve released and one local variety at Main Campus Site Hadiya zone of SNNPR. Twelve tef varieties were evaluated in RCBD with three replications on station of Main Campus Site during main cropping season of 2019/2020. Analysis of variance revealed that there were significant differences among tef varieties, Culm length, panicle length, plant height, days to heading, days to maturity, grain filling period, primary panicle brunch, grain yield, biomass yield and harvest index at Main Campus site. Based on the obtained result, the improved tef varieties namely; DZ-Cr-438 DZ-Cr-974 (Dukem), DZ-01-899 (Gimbechu) and DZ-01-196 (Magna) at Main Campus site. Therefore, these varieties showed better performance for most of the studied characters including grain yield. Therefore, these varieties were selected and recommended for the study area and similar ecologies of Hadiya Zone. This finding, being the result of one year with single location, it is recommended that the experiment should be repeated at multi locations for several years to confirm the obtained results.

## INTRODUCTION

Tef [Eragrostis tef (Zucc) Trotter] is an allotetraploid with 40 chromosomes (2n = 4x = 40), belongs to the family Poaceae, subfamily Eragrostoideae, tribe Eragrosteae and genus Eragrostis. It is an ancient and the most important cereal crop in Ethiopia and the country is considered to be both the center of origin and center of diversity for this cereal (Vavilov, 1951). There are about 350 Eragrostis species of which E.tef is the only species cultivated for human consumption. At the present time, the gene bank in Ethiopia holds over five thousand tef accessions collected from geographical regions diverse in terms of climate and elevation. Tef is adaptable to a wide range of ecological conditions in altitudes ranging from near sea level to 3000 masl and even it can be grown in an environment unfavorable for most cereal, while the best performance occurs between 1100 and 2950 masl in Ethiopia (Hailu and Seyfu, 2000). It is the major food crop in Ethiopia where it is annually cultivated on more than three million hectares of land by 6.3 million small-scale farmers on more than 30 % of the total area allocated to cereal crops (CSA, 2014). The importance of tef in Ethiopia is mainly due to its preference by both farmers and consumers. Farmers, above all, grow tef due to its tolerance to several biotic and a biotic stress especially to the poorly drained Vertisols, a dominant soil type in the central highlands where other cereals can hardly survive without the use of proper drainage system. Over 50 million people in Ethiopia consume tef as staple food due to the better quality bread called “injera” made from it compared to that from other cereals. The absence of gluten in its grain (Assefa, 2014) makes tef a healthy food such that people allergic to gluten can safely consume tef products. Compared to the straw from other cereals, the straw from tef is more nutritious and palatable for livestock feed (Kebebew et al., 2015). The nutritional value of Tef grain is similar to the traditional cereals. Tef is considered to have an excellent amino acid composition, lysine levels higher than wheat or barley, and slightly less than rice or oats. Tef is also higher in several minerals, particularly iron. Tef grain is also a rich source of protein and nutrients and has additional health benefits including that the seeds are free from gluten. According to a recent study, the bio-available iron content was significantly higher in tef bread than in wheat bread. In general, tef provides quality food and grows under marginal conditions, many of which are poorly suited to other cereals. However, tef is considered to be an orphan crop since it is only of regional importance and has until recently not been the focus of crop improvement (Assefa, 2014).

The yields of Tef are low in Ethiopia as well as in southern region due to different production problems including: lack of improved varieties, non-adoption of improved technologies, disease and pests are some of the most serious production constraints in Tef production in Ethiopia. Some varieties of Tef were released by the different regional and federal research centers in Ethiopia; however, most of them were not evaluated around areas of southern Ethiopia and farmers were not participated in varietal improvement and testing process. Participation of farmers’ in varietal choice has considerable value in technology evaluation and dissemination. Participatory varietal evaluation and selection is being conducted in some crops like common bean and barley (Fufa et al., 2010). According to Courtois et al. (2001) evaluated the effect of participation of farmers by comparing only the rankings of varieties by farmers and researchers at the same locations and reported a strong concordance between farmers and breeders in environments that have been producing contrasting plant phenotypic performance in rice. Two way feedbacks between farmers and researchers is indeed vital component of highly client-oriented breeding programs in locally important and traditionally cultivated crop (Getachew et al., 2008). Daniel et al. (2007) stated that farmers’ selection criteria vary with environmental conditions, traits of interest, ease of cultural practice, processing, use and marketability of the product, ceremonial and religious values. Therefore, this study was mainly conducted with the objectives to evaluate and select improved tef varieties which are adaptable, high yielding and to assess farmers’ criteria for variety selection with the participation of farmers in southern Ethiopia

## MATERIALS AND METHODS

### Description of the Study Area

The varieties were tested at Main Campus Agricultural Research Site (MCARS) during 2019/2020 cropping season. Main Campus Agricultural Research Site is located (7.30-7.35N, 37.48-37.52 E. It has an altitude of 2000-2500 m.a.s.l. with annual average rainfall of 850mm. The annual average temperature of the study area is16.65°C with maximum and minimum temperature of 22.8°Cand 10.5°C respectively. With the soil type classified as Nitosol soil with a pH of 5.2.

### Experimental Materials

The experimental material of the study comprised of 12 tef varieties kindly provided from the Debre Zeit Agricultural Research Center and was cultivated at Main Campus Agricultural Research Site (MCARS) during 2019/2020 cropping season.

**Table 1.** List of tef (*Eragrostis tef*) varieties used for experiments.

| No | Code | Locale name | Released By | Year of release |
| --- | --- | --- | --- | --- |
| 1 | DZ-01-196 | Magna | DZARC | 1970 |
| 2 | DZ-01-899 | Gimbechu | DZARC | 2007 |
| 3 | DZ-01-1285 | Koye | DZARC | 2002 |
| 4 | DZ-Cr-354 | Enatit | DZARC | 1970 |
| 5 | DZ-Cr-974 | Dukem | DZARC | 1995 |
| 6 | DZ-01-2675 | Degatef | DZARC | 2005 |
| 7 | DZ-Cr-438 | Kora | DZARC | 2014 |
| 8 | Ho-Cr-136 | Amarach | DZARC | 2006 |
| 9 | DZ-Cr-387 | Quncho | DZARC | 2006 |
| 10 | DZ-Cr-409 | Boset | DZARC | 2012 |
| 11 | DZ-01-255 | Gibie | DZARC | 1993 |
| 12 | DZ-CR-358 | Ziquala | DZARC | 1995 |
| 13 | Local check |  |  |  |

### Experimental Design and Trial Management

Twelve (12) improved varieties of tef were tested for their adaptability, evaluation and selection with full participation of farmers in the study areas. The trial was carried out in Randomized Complete Block Design (RCBD) in three replications. The varieties were grown under uniform rain fed conditions. The plot size was 3 m length and 3 m width (9 m^2^) with 0.2 m of row spacing. The spaces between plots and replications were 1 m and 1.5 m, respectively. Sowing was done by manual drilling along the rows at seed rate of 5 kg/ha. Sowing was done within the last week of July to 1^st^ week of August 2019. The sources of P2O5 and nitrogen fertilizer were NPS and UREA respectively. Both applied at the rate of 100 kg ha^−1^. All of the NPS was applied at planting and UREA was applied in two splits, half at the time of planting and the remaining half at tillering stage. All other pre and post-planting management practices were done in accordance with the research recommendations for tef production in the area. Twice hand weeding and plowing and other management practices were done as required. All other recommended agronomic practices were kept normal and uniform to ensure normal plant growth and development. Seed yield of each plot was recorded and then converted into kg/ha.

### Agronomic data collected

Data were collected either on plant or plot bases on yield and yield related traits.

### On plot basis

**Days to 50% heading (DH):** The numbers of days from sowing to when 50% of the plants were started heading. It was counted as the number of days from sowing to 50 % heading stage i.e., 50% of the heads fully emerged from the flag leaf sheath.
**Days to emergency**: Number of days taken from date of sowing to 50% of plants to emerge
**Days to 75% maturity (DM):** The numbers of days from date of sowing to a stage at which 75% of the plants were reached physiological maturity or 75% of the panicles on the plots turned golden yellow color.
**Grain yield per plot (GYP):** The grain yield per plot was measured in grams using sensitive balance after moisture of the seed is adjusted to 12.5%. Total dry weight of grains harvested from the middle four rows out of six rows were taken as grain yield per plot and expressed as grams per plot.
**Shoot Biomass yield per plot (BMYP):** It was recorded by weighing the total above ground yield harvested from the four central rows of each experimental plot at the time of harvest when moisture content adjusted to 8%.
**Harvest index (%):** It was estimated by dividing grain yield per plot to biological yield per plot. It is ratio of grain yield to the above ground biomass yield.

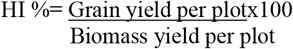

### On plant basis

**Plant height (cm):** The distance between the ground level to the tip of the terminal spikelet in cm of the mother ten plants.
**Culm length (cm)**: The heights of the ten plants selected at random were measured at harvesting time in centimeter. The height was taken as the distance between the soil surfaces to the beginning of panicle.
**Number of primary branches per plant (PPB):** Counting the total number of primary branches on main stem of each selected plant at the time of harvest
**Panicle length (cm):** Panicle lengths as the average length from the base of the panicle to the tip of ten pre-tagged plants were recorded in centimeter from central rows of each plot.
**Days to grain fill period:** Number of days from 50% heading of the plants to maturity

### Statistical analysis and variance components

The data was subjected to analysis of variance using SAS software v 9.1.3 (SAS, 2004). The Significant difference among genotypes was tested by ‘F’ test at 1% and 5% levels of Probability. The structure of analysis of variance (ANOVA) table is presented below.

**Table 2.** The structure of analysis of variance (ANOVA) (Gomez and Gomez, 1998)

| Source | Df | (SS) | (MS) | F |
| --- | --- | --- | --- | --- |
| Block | r-1 | SSB | SSB/(r-1) | MSR/MSE |
| Treatment | t-1 | SST | SST/(t-1) | MST/MSE |
| Error | (r-1)(t-1) | TSS | SSE(r-1)(t-1) |  |
|  |  | SST-SSB |  |  |
| Total | tr-1 | TSS |  |  |

Where: *r* = Number of replications; *t* = Number of treatments / genotypes; *SS* = Sum of Square; *MS* = Mean of square; *S.E.m* = $\pm E.M.SS/r$

## RESULTS AND DISCUSSION

### Analysis of Variance

The analysis of variance showed that there were highly significant (p≤0.01) difference among varieties for days to heading, plant height, grain yield, biomass yield and harvest index while significant (P≤0.05%) difference in panicle length, primary panicle brunch, Culm length, days to maturity, *days to emergency* and grain filling period at Shone site. Generally, the analysis of variance revealed that the presence of considerable variations among the 12-tef varieties for all the traits. This indicating the presence of variability, which can be exploited through selection for further breeding programs. These results were supported by Chondie, Bekele (2017) who reported considerable variation in the days to maturity, plant height and panicle length, days to heading and grain yield of different tef varieties when planted over years. Similarly, AliyiKedir *et al.* (2016) reported that highly significance differences between varieties for the characters like days to maturity, panicle length, plant height, days to heading, days to maturity and grain yield.

**Table3.** Analysis of variance for different characters of tef varieties studied at Main campus site.

| Sources of variation | df | DH | DM | DE | PL | PH | CL | PBP | GFP | GY | BM | HI |
| --- | --- | --- | --- | --- | --- | --- | --- | --- | --- | --- | --- | --- |
| MSR | 2 | 0.83 | 10.16 | 2.00 | 5.61 | 28.15 | 20.02 | 13.46 | 11.11 | 0.013 | 0.889 | 40.57 |
| MST | 16 | 12.31** | 23.52* | 3.27* | 18.2* | 33.1** | 57.3* | 45.5* | 27.8* | 0.08** | 2.45** | 130. ** |
| MSE | 32 | 3.28 | 9.64 | 1.10 | 10.01 | 20.16 | 12.16 | 7.04 | 8.23 | 0.015 | 0.5 | 24.41 |
| F-value |  | 2.06 | 2.44 | 3.00 | 1.22 | 2.6 | 4.18 | 4.69 | 2.12 | 5.67 | 4.87 | 5.33 |
| CV (%) |  | 1.78 | 2.74 | 18.30 | 8.44 | 4.09 | 5.14 | 12.1 | 8.37 | 23.26 | 23.73 | 25.73 |

\* = Significant at 5% level of probability, \*\* = highly Significant at 1% level of probability, ns= Not significant, CL=Culm length, PL=panicle length, PH=plant height, DE days to emergency, DH=days to heading, DM days to maturity, GFP=grain filling period, PPB=primary panicle brunch, GY=grain yield, BMY=biomass yield, HI=harvest index, MSR=mean square of replication, MST= mean square of treatment, CV= coefficient of variation and .

At Main Campus site, the analysis of variance indicated that there were highly significant (P<0.01) difference among varieties in plant height, days to emergency, grain yield, biomass and harvest index while significant difference (P<0.05) in Culm length, panicle length, days to heading, days to maturity and primary panicle brunch. Generally, the analysis of variance results revealed that the presence of adequate variations among the tef *varieties* for most of the traits suggesting that the higher opportunity of selecting *varieties* for trait of interest. The result of analysis of variance allows carrying out further genetic analyses for all traits. Similarly, Asaye 2017 reported that days to heading, days to maturity, grain yield per plot and shoot biomass. Grain filling period is only character that show non significance. Likewise, Hailu *et al.* (2003a) reported significant differences in days to heading, days to maturity, plant height, panicle length, panicle weight, yield per panicle, lodging index, shoot biomass and grain yield.

### Range and Mean Values

The mean performances of the Twelve Tef varieties and one Local checks for 11 characters are presented in Table 5. The mean values for days to 75% maturity ranged from 117.5 (DZ-Cr-354) to 108.2.75(DZ-01-1285), Plant height was varies from 94.0 (DZ-Cr-438) to 81.2 (DZ-Cr-354), The mean values for days to 50% heading ranged from 74.5 (DZ-Cr-387) to 67.0 (Ho-Cr-136), Culm length was ranged from 67.1 (DZ-Cr-974) to 49.7 (DZ-Cr-354), palm length was ranged from 36.4 (DZ-Cr-438) to 24.7 (DZ-Cr-974), Number of primary branches per plant was ranged from 25.2 (DZ-Cr-974) to 14.7 (DZ-01-2675), Grain filling is an important trait that ultimately affects the overall grain yield by increasing grain weight. Therefore, it was ranged from 45.2(DZ-Cr-354) to 35.2 (DZ-01-1285), Days to emergency was ranged from 7.0 (DZ-Cr-387) to 3.56 (Ho-Cr-136). Grain yield was ranged from 1955 (DZ-Cr-974) to 490 (DZ-01-1285), biomass yield per plot and harvest index was ranged from 5.1 (DZ-Cr-974) to 2 (DZ-01-2675), 33.2 (Ho-Cr-136) and 6.4 (DZ-01-1285) respectively. From the result it was observed that those characters with the higher range of values were also had higher mean values and vice versa. Such considerable range of variations provided a good opportunity for yield improvement. Thus, high variability for 11 traits in twelve and one local check studies implied that there was reasonably sufficient variability. This provides ample scope for selecting superior and desired Tef varieties by the plant breeders for further improvement. Generally, the range of variation was wide for all the characters studied at Main Campus site. Abebe (2013) also reported wide range of variation among Tef genotypes. But this result is in contrast to Fentie *et al.,* (2012) finding.

The mean performances of the Twelve Tef varieties and one Local checks for 11 characters are presented in Table 6. The mean values for Grain yield was ranged from 1723 (DZ-Cr-438) to 950 (DZ-Cr-354), days to 75% maturity ranged from 105 (Local check) to 98.2 (DZ-01-1285), Plant height was varies from 103.2 (DZ-Cr-438) to 78.2 (DZ-01-196), The mean values for days to 50% heading ranged from 72.3 (Local check) to 62.4 (DZ-Cr-409), Culm length was ranged from 67.6 (DZ-Cr-438) to 45.5 (DZ-01-196), Grain filling is an important trait that ultimately affects the overall grain yield by increasing grain weight. Therefore, it was ranged from 36.5 (DZ-Cr-354) to 25.5 (DZ-Cr-438), palm length was ranged from 37.2 (DZ-CR-358) to 25.9 (DZ-Cr-974), Number of primary branches per plant was ranged from26.8 (DZ-CR-358) to 16.0 (DZ-01-899), Days to emergency was ranged from 8.0 (DZ-Cr-387) to 3.5 (DZ-01-196). Biomass yield per plot and harvest index was ranged from 3.8 (DZ-Cr-438) to 1.73 (DZ-Cr-387), 26.4 (DZ-Cr-387) and 15.3 (DZ-Cr-409) respectively. From the result it was observed that those characters with the higher range of values were also had higher mean values and vice versa. Such considerable range of variations provided a good opportunity for yield improvement. Thus, high variability for 11 traits in twelve and one local check studies implied that there was reasonably sufficient variability. This provides ample scope for selecting superior and desired Tef varieties by the plant breeders for further improvement. Generally, the range of variation was wide for all the characters studied at Main Campus site. Similar results were previously reported by Abebe, 2013.

**Table 6.**
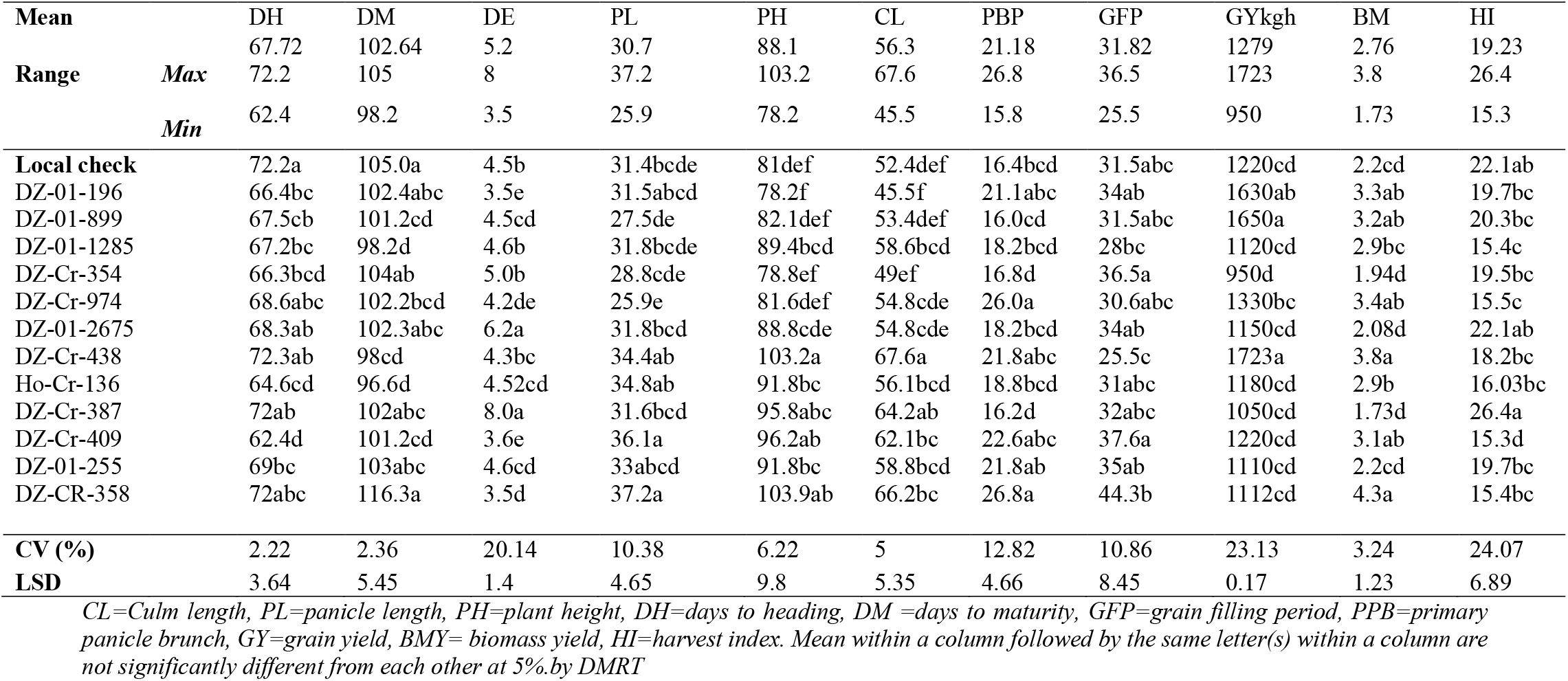
Mean and Range values for different agronomic traits for 12 Tef varieties at Main Campus site main cropping season 2019/2020.

## CONCLUSION AND RECOMMENDATION

The objective of present investigation was to evaluate and select improved tef varieties which are adaptable, high yielding and to assess farmers’ criteria for variety selection with the participation of farmers. Analysis of variance means performance of quantitative traits in this study showed that there were significant differences among tef varieties for Grain yield, days to maturity, plant height, days to heading, Culm length, Grain filling period, palm length, Number of primary branches per plant, Days to emergency, biomass yield and harvest index at Main Campus site. High grain yield of tested varieties at Main Campus site recorded by variety DZ-Cr-438-(1723kg/h) followed by DZ-Cr-354 (950 kg/h). On the other hand, lowest grain yield was recorded by DZ-Cr-354 (950 kg/h). Grain yield was an important character to be considered for variety selection to address the objective of the present activity. For this reason, three improved varieties i.e. DZ-Cr-438 DZ-Cr-974 (Dukem), DZ-01-899 (Gimbechu) and DZ-01-196 (Magna) at Main Campus site. Therefore, these varieties were selected and recommended for the study area and similar ecologies of Hadiya Zone and being the result of one year with single location; it is recommended that the experiment should be repeated at multi locations for several years to confirm the obtained results.

## ACKNOWLEDGEMENTS

We would like to express our sincere gratitude to Wachemo University, for granting full fund of the project and Wachemo University Research Development Directorate Office for facilitating the process and services required for the completion of the research projects. My special thanks go to Wachemo University, Collage of Agricultural science, Department of Plant science and Field Assistants for their contribution to facilitate field operation and other activities. We would like to extend our heartfelt gratitude to Wachemo University finance office for their guidance to the budget allotted to the project as per the proposal respecting the financial rules and regulation and Debrezeyt Agriculture Research Center for contribution of seed.

